# ProbAlign: a re-alignment method for long sequencing reads

**DOI:** 10.1101/008698

**Authors:** Feng Zeng, Rui Jiang, Guoli Ji, Ting Chen

## Abstract

The incorrect alignments are a severe problem in variant calling, and remain as a challenge computational issue in Bioinformatics field. Although there have been some methods utilizing the re-alignment approach to tackle the misalignments, a standalone re-alignment tool for long sequencing reads is lacking. Hence, we present a standalone tool to correct the misalignments, called ProbAlign. It can be integrated into the pipelines of not only variant calling but also other genomic applications. We demonstrate the use of re-alignment in two diverse and important genomics fields: variant calling and viral quasispecies reconstruction. First, variant calling results in the Pacific Biosciences SMRT re-sequencing data of NA12878 show that false positives can be reduced by 43.5%, and true positives can be increased by 24.8% averagely, after re-alignment. Second, results in reconstructing a 5-virus-mix show that the viral population can be completely unraveled, and also the estimation of quasispecies frequencies has been improved, after re-alignment. ProbAlign is freely available in the PyroTools toolkit (https://github.com/homopolymer/PyroTools).

## Introduction

The advent of high throughput sequencing (HTS) technologies has revolutionized the discovery of genetic changes in genome sequence. Various sizes of samples, such as a set of targeted regions in a genome, or the entire genome of a sample, even the genomes of a pool of samples, can be completely sequenced in days or weeks, producing huge data of gigabase-size. In a genome, genetic variants, e.g. single nucleotide polymorphisms (SNPs), multiple nucleotide polymorphisms (MNPs), short insertions and deletions (InDels), can be pinpointed at the base pair (bp) resolution by using the re-sequencing or targeted sequencing technique. Leveraging on the multiplex indexing technique, a population of tens of samples can be sequenced by using the same sequencing runs and completed in feasible time with affordable cost. Population sequencing data gears up the genotyping of genetic variants in a population, and expedites the estimation of the distribution of genetic variants in the population, which eventually facilitating the dissection of population structure [1], the sketching of fitness landscape [2], the reconstruction of the clonal evolution [3], etc.

There are four kinds of sequencers that dominate the market, including Illumina [4], Roche/454 [5], Ion Torrent [6], and Pacific Biosciences SMRT [7]. While all of these sequencers can produce a huge amount of sequencing reads, they are different at various aspects, e.g. reagent cost per base, data throughput, read length, error patterns, etc. The Illumina sequencer is commonly used in the large-scale genomics projects that require the sequencing of a large population or high sequencing depth, because it achieves the lowest per base sequencing cost and the largest data throughput. The length of Illumina sequencing reads is within the range from 100bp to 300bp. Although this interval covers most of microsatellite repeats and short interspersed nuclear elements (SINEs), short reads limit the ability of resolving the complex regions in genome, e.g. long interspersed nuclear elements (LINEs), of which the size is from 500bp to 8000bp [8]. Roche/454 and Ion Torrent sequencers can produce the mediate length of sequencing reads ranging from 300 bp to 800 bp. Pacific Biosciences SMRT sequencer, which using third-generation single molecule sequencing technology, can produce long reads up to 10kilo base pairs. Mediate and long reads are less sensitive to the repetitive regions, and then achieve the better mappability than short reads, so as to move forward the reconstruction of complex regions in genome [9]. Besides the discrepancy at read length, the error patterns of these sequencers are different. The Illumina sequencer is prone to substitute the underlying bases with erroneous bases [10]. In Roche/454 and Ion Torrent data, the insertions and deletions are prevalent because they are imprecise at the determination of the length of homopolymer runs [11]. The insertions and deletions are also the dominant error patterns in the Pacific Biosciences SMRT data [12]. All of these four kinds of sequencers play the important roles in the genomic research fields.

Variant calling is the first step of HTS applications, and proceeds in the following stages. First, sequencing reads are mapped onto the reference genome, and piled up on the genome. Second, the differences are detected by comparing sequencing reads with the genome. Among the called variants, the false discoveries are resulted from three confounding factors: sequencing errors, alignment biases, and mapping artifacts. (i) Base quality score and error modeling are employed in both Likelihood- and Bayesian-based variant calling methods in order to recognize the truth genetic signals and eliminate sequencing errors [13–15]. (ii) The gaps in the short tandem repeats, e.g. homopolymer runs and dinucleotide repeats, are usually placed at wrong positions, then resulting in the incorrect alignments and also the false positive variants. It is a challenge to compute the correct alignments in these low complexity regions, because (a) sequencing reads vary in a large range at determining the number of repetitive elements, and (b) there is no any proper scoring function, which reflecting error patterns of a sequencer. Alignment biases are pervasive in the mediate and long sequencing reads. (iii) The repetitive elements in genome, e.g. satellites, rDNAs, SINEs, LINEs, *Alu* repeats, segmental duplications, etc., place a problem at searching for the original positions of sequencing reads, especially for short reads. A short read is often randomly assigned to a position by a mapping algorithm, if it could be mapped to multiple regions of the similar repetitive structure. The differences between the repetitive regions could be reported as genetic variants, eventually increasing the false discovery rate. Moreover, the alignment biases could place the truth variants at random positions, which decreasing the number of the variant supporting reads in the pileup, and then result in the false negatives. The mapping artifacts that place reads in wrong positions are also a resource of the false negatives.

Haplotype-based variant calling has been reported as an efficient variant calling method [16–19]. The computation steps include: (1) generating the candidate haplotypes, (2) computing the new alignments between the sequencing reads and candidate haplotypes, which is also called re-alignment, and (3) inferring the underlying genotype. The improvement of haplotype-based methods in variant calling relies on the following advantages. (i) First, haplotype-based methods employ the graph technique, e.g. de Bruijn graph [20] or partial order graph [21], to reconstruct the consensus sequences (haplotypes) within a region of interest (ROI). The strength of the graph technique is to remove the spurious signals by using the redundancy in the pileup of sequencing reads. The underlying genotype is a combination of the consensus sequences, which can explain the largest portion of the aligned reads in the ROI region. As a consequence of the mapping artifacts, the number of unique reads in a ROI region could be more than the expected number of haplotypes, e.g. two haplotypes for a diploid sample. The genotype inference places the constraints on the number of haplotypes, and then eliminates the mapping artifacts. (ii) Second, haplotype-based methods approximate the truth alignments of the sequencing reads by enumerating over all possible haplotypes. The misalignments can be corrected if sequencing reads were aligned to their original haplotypes.

However, the re-alignment is commonly implemented as an internal workhorse inside the haplotype-based variant calling methods, and it is not able to access to the re-alignment results from the exterior. GATK provides a tool, which called IndelRealigner, to adjust the alignments around the insertions/deletions, but it is restricted to process the Illumina sequencing data [14]. SRMA is another re-alignment tool, but its speed is a critical issue and cannot work on Pacific Biosciences SMRT data [22]. To our knowledge, there is no any standalone re-alignment tool that working for long read sequencers under the feasible running time in the Bioinformatics field. The high quality alignments can also benefit other genomic applications, e.g. the detection of short tandem-repeat variation [23] and viral quasispecies reconstruction [24]. There are two difficulties in the implementation of the re-alignment. First, the re-alignment is a time-consumed computation, because it requires the computation of the pairwise alignments between sequencing reads and all possible haplotypes. Second, there is no any proper scoring function that can correctly score the alignments for sequencing reads, of which sequencing errors vary in a diverse range.

In this paper, we presented a standalone re-alignment tool, called ProbAlign, for long read sequencing technologies. ProbAlign addressed the above two difficulties. First, ProbAlign employs the *graph* technique to represent the *consensus sequences* in a ROI region, which resembles the partial order graph or variant graph. ProbAlign computes the new alignment of a sequencing read by aligning the read to the graph. The time complexity is *O*(*n*|*V*|), *n* is read length, |*V*| is the number of vertices in the graph. As a comparison, the time complexity of the re-alignment in the haplotype-based methods is *O*(*λmn*), *λ* is the number of haplotypes, *m* is haplotype length, and *n* is read length. Hence, the time complexity is reduced by an order of magnitude. Second, ProbAlign employs the *conditional random field* (CRF) technique to model error patterns in the sequencing reads. The parameters of the CRF model can be estimated from the BAM file at hand. This enables the proper scoring of the correct alignments for sequencing reads, which are contaminated by various sequencing errors.

## Results

### Overview of ProbAlign

The rationale of the re-alignment method is stated in the following. Assume that a bucket of reads have been aligned into a multiple sequence alignment (MSA), of which a row is an aligned read and a column is an aligned position. The computational task is to align a new read onto the MSA. The strategy, which implemented in ProbAlign, is to pick up a sequence in the MSA, and then align the new read to the picked sequence. The sequence is picked such that sequencing errors in the sequence are most similar to those in the new read.

As shown in Fig. 1, ProbAlign defines low complexity regions, e.g. homopolymer runs and dinucleotide repeats, and putative multiple nucleotide polymorphism (MNP) loci as the regions of interest (ROIs). (i) First, a DAG is established from the raw mapping results in a ROI. In the DAG, A vertex represents one allele in an aligned position. An edge links two vertices that are occurred in an aligned read. (ii) Second, the vertices in the DAG are sorted in the topological order, and a depth-first search (DFS) algorithm is deployed to find the maximally weighted path(s) in the graph, which are inferred as the consensus sequence(s) in the ROI. (iii) Next, a new DAG is constructed from the reference sequence and the consensus sequence(s). This DAG is named as *the consensus graph*, because every path is a consensus sequence. (iv) Then, ProbAlign iterates through the sequencing reads in the ROI, and the alignments of sequencing reads against the consensus graph are computed by using a Viterbi algorithm. Once the new alignment of a sequencing read is better than the original alignment, e.g. the number of mismatches declines, and then the new alignment replaces the old alignment in the given BAM file. Also, the consensus graph is updated by adding new vertices and/or edges, which representing the variants in the new alignment. The reason of the graph update is that the number of short repetitive elements varies in a diverse range in long sequencing reads, the alignment quality can be improved if the graph has the local structures describing the large deletions/insertions in low complexity regions. (v) Finally, the CIGAR strings and MD tags of the re-alignments are stored in the new BAM file. Two examples of the re-alignment are shown in Fig. 2. More re-alignment examples could be found in the supplemental materials.

**Figure 1.**
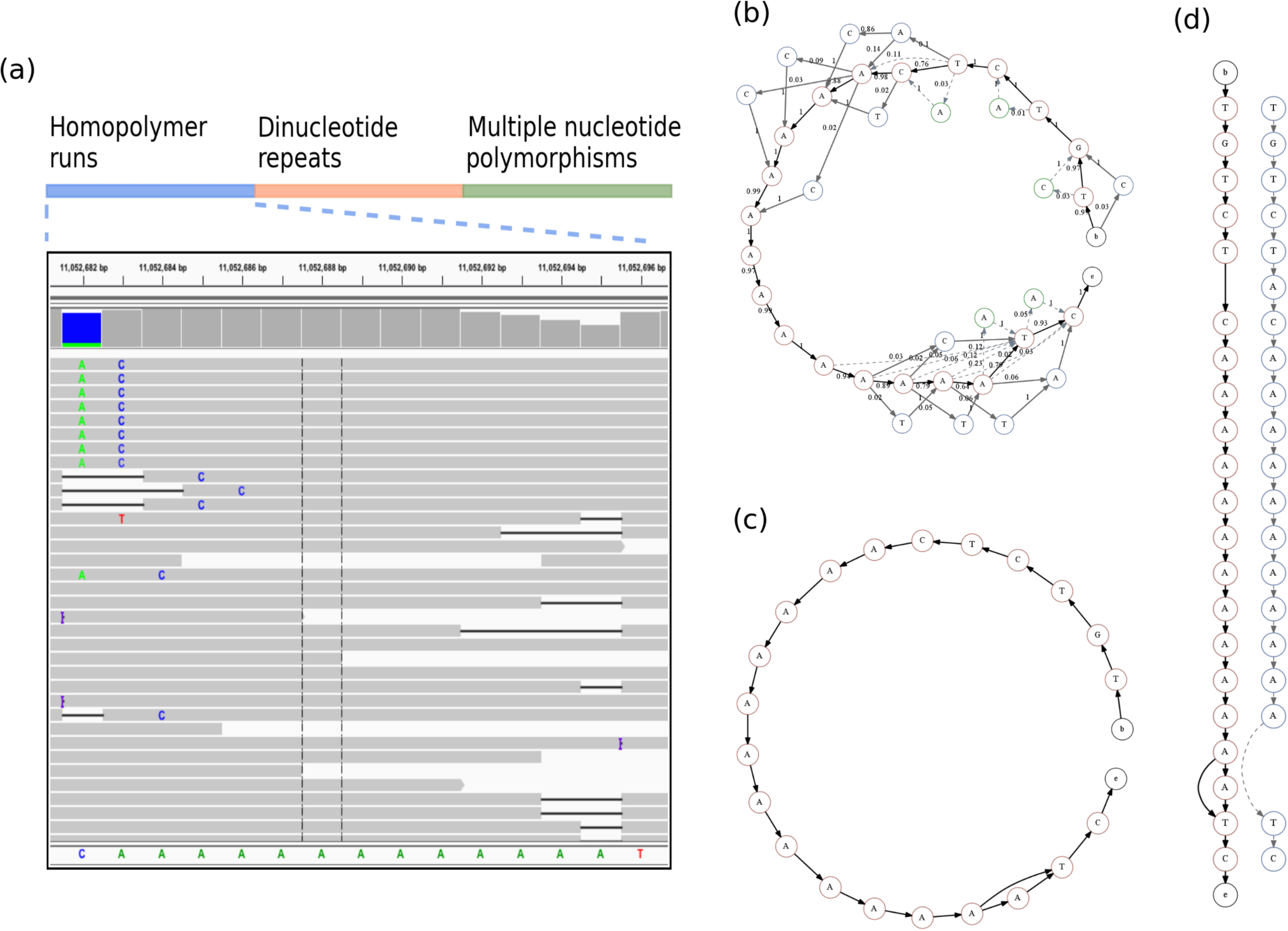
Workflow of ProbAlign. ProbAlign adjusts the misalignments around homopolymer runs, dinucleotide repeats, and multiple nucleotide polymorphisms. (a) A region of interest includes a homopolymer run of length 13 bp, and also harbors a number of the incorrect alignments. (b) An alignment graph is constructed from the pileup. Red vertices in the backbone are the bases in reference genome. Blue vertices are the mismatches. Green vertices are the inserted bases. Dashed edges represent insertions and deletions. Weight of an edge *e* = 〈*u,v*〉 is computed by the formula, 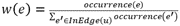. (c) A consensus graph is constructed from the alignment graph. (d) It is an alignment of a read sequence (colored blue) against the consensus graph (colored red).

**Figure 2.**
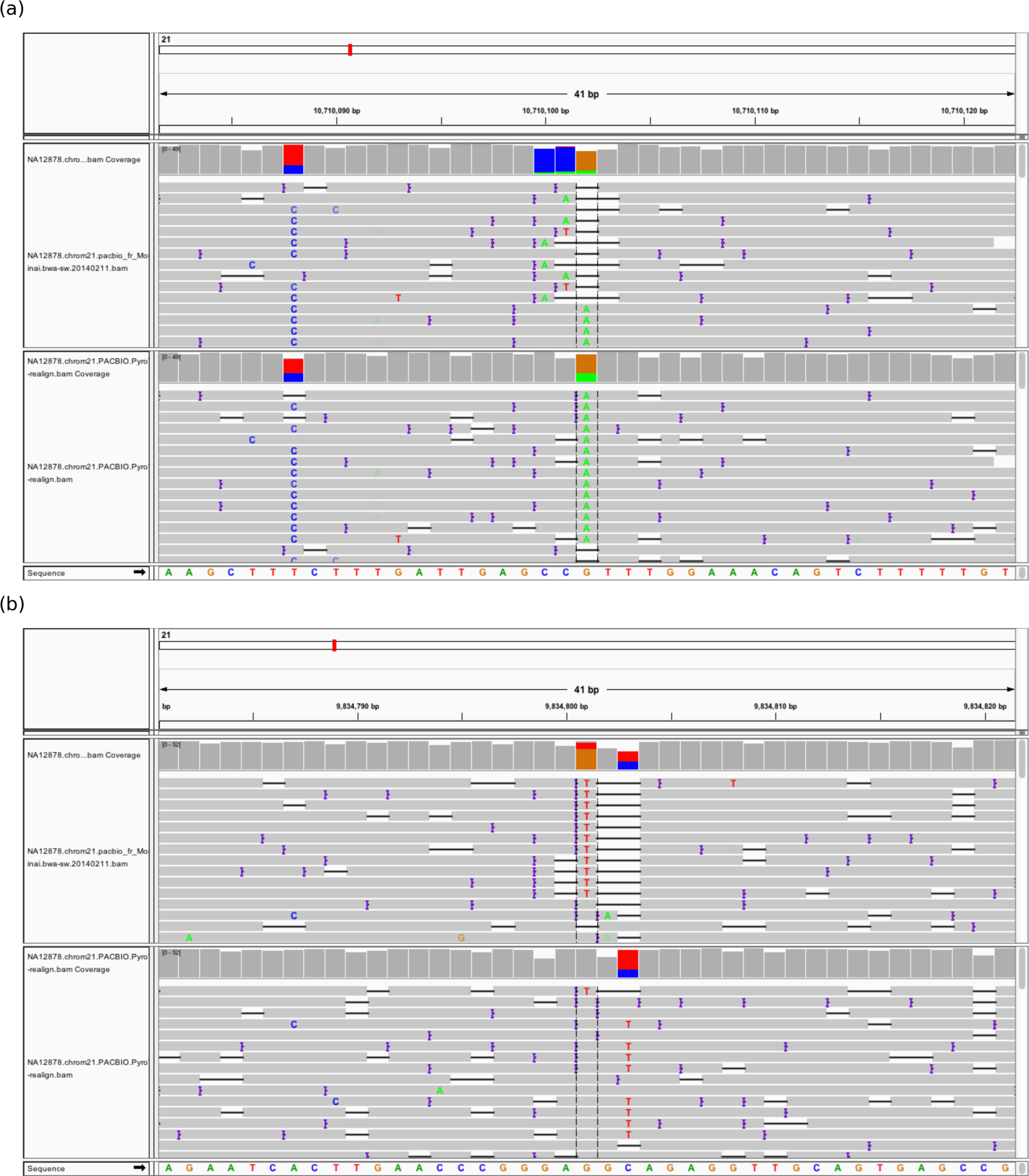
Re-alignment examples of long sequencing reads. The tracks from top to bottom are the original BAM file, the re-aligned BAM file, and human genome. (a) An example of re-alignment increasing true positive detection. The truth SNP at chr21:10710102 is rescured by replacing the misalignments. (b) An example of re-alignment reducing false positive detection. The false SNP at chr21:9834810 is reduced by replacing the misalignments.

### Application I: SNP calling of NA12878

The alignment files of Pacific Biosciences SMRT re-sequencing data (unpublished) of NA12878 that sequenced by Mount Sinai School of Medicine are publicly available in the 1000 Genomes Project FTP site. The alignment files of Roche/454 re-sequencing data of NA12878 was also downloaded from the 1000 Genomes Project FTP site. The Ion Torrent re-sequencing reads of NA12878 were downloaded from the National Center for Biotechnology Informatics (NCBI) Short Read Archive (SRA) (Project Accession Number: PRJNA162355). The Ion Torrent sequencing reads were mapped to the human genome (version: hg19) by using the Burrows-Wheeler Aligner (BWA) MEM algorithm [25] with the default option settings.

The alignments in the BAM files of Roche/454, Ion Torrent, and Pacific Biosciences SMRT reads were adjusted by using ProbAlign in the default settings. The SNP calling software Samtools [13] was deployed to call SNPs in both the original and adjusted BAM files. The VCF files of SNP calling results were compared with the National Institute of Standards and Technology (NIST) GIAB high- confidence benchmark calls (v2.18), which constructed by the Genome in a Bottle (GIAB) consortium by integrating the variant calls from five sequencers, including Illumina, Roche/454, Ion Torrent, SOLiD and Complete Genomics [26]. The evaluation measures, e.g. the number of true positives, the number of false positives, and the false discovery rate (FDR), were calculated by using the program VCFComparator in USeq toolkit.

Taking Pacific Biosciences SMRT data for example, the number of sequencing reads that mapped to chromosome 21 is 661553, of which the average read length is 1805bp. ProbAlign found 70581 ROIs in chromosome 21, and re-aligned 244986 (37.0%) sequencing reads. Both the original BAM file and re-aligned BAM file were input to Samtools to call SNPs by using the same options. When the cutting threshold of variant posterior probability was set as 0.5, which implying that variant quality score should be greater than 3.02, Samtools called 41348 true positives and 1815 false positives in the re-aligned BAM file. As a comparison, The Samtools caller detected 41322 true positives and 3214 false positives in the original BAM file. As a result of re-alignment, the true positives were increased by an amount of 26 (0.1%), and the number of true negatives was decreased by 1399 (43.5%). We plotted the increment of true positives in Fig. 3a, by varying the cutting threshold of variant quality score. It is noted that the re-alignment improved the probability of truth SNPs being detected, especially increasing the number of high quality truth SNPs. As shown in Fig. 3a, the average relative increment percentage reaches 24.8%. We also plotted the decrement of false positives along with the variant quality score in Fig. 2b. It revealed that the re-alignment efficiently suppress the false positive detection. The re-alignment method helps decrease a considerable number of false positives in the low quality SNPs, e.g. over 40% of false positives have been eliminated at the variant quality score threshold of 10. Although the figure of the relative decrement percentage shows that the re-alignment method harasses the detection of false positives in the high quality calls, the absolute number is below 5 and could be overlooked. Generally speaking, the re-alignment decreases the false discovery rate (Fig. 3c), as well as increases the capability at detecting high quality variants. In both Roche/454 and Ion Torrent data, the false discovery rates were also reduced. Those results can be found in the supplemental materials.

**Figure 3.**
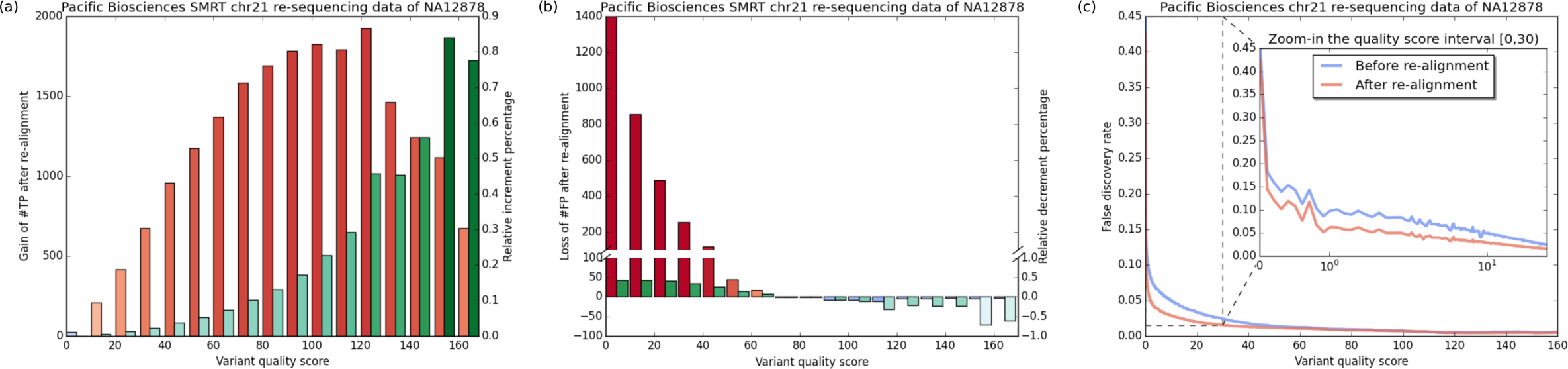
The increment of true positives, the decrement of false positives, and FDR curve. (a) The increment of true positives, red bars are the absolute changes, and green bars are the relative changes. (b) The decrement of false positives, red bars are the absolute changes, and green bars are the relative changes. (c) FDR curve.

### Application II: HIV-1 quasispecies reconstruction

We also evaluated the performance of ProbAlign in viral population analysis. We used a public HIV-1 benchmark data including five HIV-1 strains, which are HIV-1_89.6_, HIV-1_HXB2_, HIV-1_JR-CSF_, HIV- 1_NL43_, and HIV-1_YU2_ [27]. The proportion of these five strains in the mix was measured by the data provider who using the single-genome amplification (SGA) method. The SGA measures are used as the golden standard to evaluate the goodness of HIV-1 quasispecies reconstruction. We used the BWA MEM algorithm to map the Pacific Biosciences SMRT sequencing data onto the genome sequence of HIV_HXB2_. The Pacific Biosciences SMRT sequencing data harbors unexpected high rate of the indels, such that it is difficult to reconstruct the quasispeices in the mix. Hence, we deployed ProbAlign to adjust the alignments in the region of the gene *gp41* that starting from 7758 to 8795 in the HIV_HXB2_ genome. We utilized the haplotype inference program PredictHaplo [28] to reconstruct the quasispecies in this region. As shown in Table 1, PredictHaplo reconstructed 4 quasispecies, if the alignments were not adjusted. The strain HIV_HXB2_ is missed in the reconstructed population. After using ProbAlign to adjust the alignments, PredictHaplo successfully reconstructed all sequences of those five HIV-1 strains. Also, the re-alignment improved the estimation of viral quasispecies frequencies. After re-alignment, the root-mean- square error (RMSE) of frequency estimation is reduced from 0.101 to 0.058. Therefore, besides the SNP calling, the re-alignment also helps resolving the structure of viral population.

**Table 1.** Reconstruction of HIV gene *gp41* in 5-virus-mix

| HIV-1 strains | Estimated quasispecies frequencies% |  |  |  |  |
| --- | --- | --- | --- | --- | --- |
|  | 89.6 | HXB2 | JR-CSF | NL43 | YU2 |
| Golden standard | 12.3 | 9.2 | 27.9 | 38.4 | 12.2 |
| Before re-alignment | 14.7 | na | 23.6 | 57.0 | 4.7 |
| After re-alignment | 14.6 | 8.7 | 23.2 | 48.1 | 5.4 |

## Discussions

*Where to re-alignment?*
*What difficulties remain in variant calling?*
How to set the scoring function?
*Computational time*
Conclusions

## Methods

### Conditional random field

Conditional random field (CRF) is a machine learning method for the processing and manipulation of sequential data, e.g. next-generation sequencing reads. CRF, like hidden Markov model (HMM), can compute the alignment of two sequences by using the *Viterbi* algorithm [29]. And also, CRF provides more flexible utilities at modeling sequence context in the alignment. Three kinds of distinct feature functions are employed in the proposed CRF model. The feature functions are defined as *f*(**x**,**y**,**a*_i,j_***), of which ***a**_i,j_* is the alignment of two subsequences **x**_[1:*i*]_ and **y**_[1:*j*]_ If a feature appears in the alignment, then *f*(**x**,**y**,**a*_i,j_***) = 1. Otherwise, *f*(**x**,**y**,**a*_i,j_***) = 0. (1) First, one kind of these feature functions is to describe the hidden state transition between two contiguous positions, e.g. previous state is *match* and current state is *insert*. (2) The second type of feature functions is the nucleotide emission in the alignment, e.g. **x***_i_* = *A* emits **y***_j_* = *A* at the hidden state *match*. (3) The rest of feature functions are designed to formulate the homopolymer insertions/deletions in the alignment, e.g. the largest length is 5 bp and the difference is 1 bp in the alignment of **x**_[*i*−4:*i*]_ = *AAAA* to **y**_[*j*−5:*j*]_ = *AAAA* at the hidden state *insert*.

There are 76 feature functions in the CRF model in total, 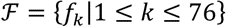. A weight *w_k_* is assigned to a feature function *f_k_* to reflect the importance of the feature. Then, the scoring of an alignment **a** is

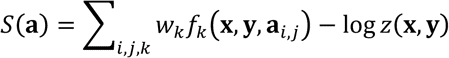

The partition function 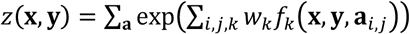 is computed by using the *forward* algorithm. The value of feature function weights *w*_*k*_ are estimated by using the iterative learning method and quasi-Newton optimum searching technique [30].

### Graph based re-alignment

The graph based multiple sequence alignment (MSA) was early proposed to resolve the gap representation problem of the consensus/profile based MSA that usually charges the gap incorrectly [21]. Recently, the graph based MSA technique has been re-invented to reconstruct the consensus sequence(s) from high throughput sequencing data, because the graph provides the structure to represent the linkage information between nearby variants, and is capable to eliminate random errors in the noise-contaminated sequencing data. This idea was employed in recent developed Bioinformatics tools, which are used to detect variants [18] or assemble long reads of high error rate [31]. The graph technique was also employed to adjust the alignments around the insertions/deletions, by leveraging on the consensus structure of the pileup of aligned reads [22]. However, the scoring function is a critical issue that the trivial scoring function cannot resolve sequencing errors in complex regions, e.g. the tandem repetitive regions. Therefore, a contribution of this paper is to propose a machine learning method to estimate the scoring function from high-throughput sequencing data. Another contribution is to employ the graph- based MSA to resolve the misalignments around tandem repeats, e.g. mono- and di-nucleotide repeats.

The re-alignment method is stated in the following steps. (1) The re-alignment method starts from the raw alignments that are generated by a mapping program. A graph is constructed from the raw alignments, of which the vertices are nucleotides in the alignments. Two vertices are linked by a directed edge if these two adjoining nucleotides occur in an alignment. The spurious edges (e.g. the edge occurs once in data) are pruned away. If the raw alignments were consistent, the re-alignment method would halt. Otherwise, the re-alignment method will proceed. The raw alignments are *inconsistent* if there are at least two paths in the graph, of which the path labels are the same. (2) Once the raw alignments are inconsistent, the top ranked *k* paths are chosen as the possible underlying consensus sequences, and the rest of paths are removed from the graph, excluding the reference. Then, all vertices in the graph are sorted by topological ordering. (3) A read is aligned against the graph by using the *Viterbi* algorithm. After the computation, a path in the graph is chosen as the aligned template, and the new CIGAR flag of the alignment between the read and template is reported. And then, the graph is updated by the computed alignment. This step is repeated until that all reads are aligned to the graph.

### Datasets and other programs

Pacific Biosciences SMRT sequencing data of NA12878 ftp://ftp.1000genomes.ebi.ac.uk/vol1/ftp/technical/working/20131209_na12878_pacbio/Schadt/alignment/

Roche/454 sequencing data of NA12878 http://ftp.1000genomes.ebi.ac.uk/vol1/ftp/technical/pilot2_high_cov_GRCh37_bams/data/NA12878/alignment/

GIAB high-confident benchmark calls of NA12878 ftp://ftp-trace.ncbi.nih.gov/giab/ftp/data/NA12878/variant_calls/

5-virus-mix https://github.com/armintoepfer/5-virus-mix

USeq toolkit http://sourceforge.net/projects/useq/

PredictHaplo http://bmda.cs.unibas.ch/HivHaploTyper/

## Reference

1. Nair S, Nkhoma SC, Serre D, Zimmerman PA, Gorena K, Daniel BJ, Nosten F, Anderson TJC, Cheeseman IH: Single-cell genomics for dissection of complex malaria infections. Genome Res 2014, **24**:1028–1038.

2. Acevedo A, Brodsky L, Andino R: Mutational and fitness landscapes of an RNA virus revealed through population sequencing. Nature 2013, advance on.

3. Wang Y, Waters J, Leung ML, Unruh A, Roh W, Shi X, Chen K, Scheet P, Vattathil S, Liang H, Multani A, Zhang H, Zhao R, Michor F, Meric-Bernstam F, Navin NE: Clonal evolution in breast cancer revealed by single nucleus genome sequencing. Nature 2014, 512:155–160.

4. Bentley DR, Balasubramanian S, Swerdlow HP, Smith GP, Milton J, Brown CG, Hall KP, Evers DJ, Barnes CL, Bignell HR, Boutell JM, Bryant J, Carter RJ, Keira Cheetham R, Cox AJ, Ellis DJ, Flatbush MR, Gormley NA, Humphray SJ, Irving LJ, Karbelashvili MS, Kirk SM, Li H, Liu X, Maisinger KS, Murray LJ, Obradovic B, Ost T, Parkinson ML, Pratt MR, et al.: Accurate whole human genome sequencing using reversible terminator chemistry. Nature 2008, 456:53–9.

5. Margulies M, Egholm M, Altman WE, Attiya S, Bader JS, Bemben LA, Berka J, Braverman MS, Chen Y-J, Chen Z, Dewell SB, Du L, Fierro JM, Gomes X V, Godwin BC, He W, Helgesen S, Ho CH, Ho CH, Irzyk GP, Jando SC, Alenquer MLI, Jarvie TP, Jirage KB, Kim J-B, Knight JR, Lanza JR, Leamon JH, Lefkowitz SM, Lei M, et al.: Genome sequencing in microfabricated high-density picolitre reactors. Nature 2005, 437:376–80.

6. Rothberg JM, Hinz W, Rearick TM, Schultz J, Mileski W, Davey M, Leamon JH, Johnson K, Milgrew MJ, Edwards M, Hoon J, Simons JF, Marran D, Myers JW, Davidson JF, Branting A, Nobile JR, Puc BP, Light D, Clark TA, Huber M, Branciforte JT, Stoner IB, Cawley SE, Lyons M, Fu Y, Homer N, Sedova M, Miao X, Reed B, et al.: An integrated semiconductor device enabling non-optical genome sequencing. Nature 2011, 475:348–52.

7. Eid J, Fehr A, Gray J, Luong K, Lyle J, Otto G, Peluso P, Rank D, Baybayan P, Bettman B, Bibillo A, Bjornson K, Chaudhuri B, Christians F, Cicero R, Clark S, Dalal R, Dewinter A, Dixon J, Foquet M, Gaertner A, Hardenbol P, Heiner C, Hester K, Holden D, Kearns G, Kong X, Kuse R, Lacroix Y, Lin S, et al.: Real-time DNA sequencing from single polymerase molecules. Science 2009, 323:133–8.

8. Treangen TJ, Salzberg SL: Repetitive DNA and next-generation sequencing: computational challenges and solutions. Nat Rev Genet 2012, 13:36–46.

9. Huddleston J, Ranade S, Malig M, Antonacci F, Chaisson M, Hon L, Sudmant PH, Graves TA, Alkan C, Dennis MY, Wilson RK, Turner SW, Korlach J, Eichler EE: Reconstructing complex regions of genomes using long-read sequencing technology. Genome Res 2014:gr.168450.113–.

10. Minoche AE, Dohm JC, Himmelbauer H: Evaluation of genomic high-throughput sequencing data generated on Illumina HiSeq and genome analyzer systems. Genome Biol 2011, 12:R112.

11. Huse S, Huber J, Morrison H: Accuracy and quality of massively parallel DNA pyrosequencing. Genome Biol 2007, 8:R143.

12. Ross MG, Russ C, Costello M, Hollinger A, Lennon NJ, Hegarty R, Nusbaum C, Jaffe DB: Characterizing and measuring bias in sequence data. Genome Biol 2013, 14:R51.

13. Li H, Ruan J, Durbin R: Mapping short DNA sequencing reads and calling variants using mapping quality scores. Genome Res 2008, 18:1851–8.

14. DePristo MA, Banks E, Poplin R, Garimella KV, Maguire JR, Hartl C, Philippakis AA, del Angel S, Rivas MA, Hanna M, McKenna A, Fennell TJ, Kernytsky AM, Sivachenko AY, Cibulskis K, Gabriel SB, Altshuler D, Daly MJ: A framework for variation discovery and genotyping using next. generation DNA sequencing data. Nat Genet 2011, 43:491–8.

15. Zeng F, Jiang R, Chen T: PyroHMMsnp: an SNP caller for Ion Torrent and 454 sequencing data. Nucleic Acids Res 2013, 41:e136.

16. Albers CA, Lunter G, MacArthur DG, McVean G, Ouwehand WH, Durbin R: Dindel: accurate indel calls from short-read data. Genome Res 2011, 21:961–73.

17. Garrison E, Marth G: Haplotype-based variant detection from short-read sequencing. arXiv:12073907 2012:9.

18. Zeng F, Jiang R, Chen T: PyroHMMvar: a sensitive and accurate method to call short indels and SNPs for Ion Torrent and 454 data. Bioinformatics 2013, 29:2859–68.

19. Rimmer A, Phan H, Mathieson I, Iqbal Z, Twigg SRF, Wilkie AOM, McVean G, Lunter G: Integrating mapping., assembly. and haplotype-based approaches for calling variants in clinical sequencing applications. Nat Genet 2014, 46:912–918.

20. Pevzner PA, Tang H, Waterman MS: An Eulerian path approach to DNA fragment assembly. Proc Natl Acad Sci U S A 2001, 98:9748–53.

21. Lee C, Grasso C, Sharlow MF: Multiple sequence alignment using partial order graphs. Bioinformatics 2002, 18:452–464.

22. Homer N, Nelson SF: Improved variant discovery through local re-alignment of short. read next-generation sequencing data using SRMA. Genome Biol 2010, 11:R99.

23. Press M, Carlson KD, Queitsch C: The overdue promise of short tandem repeat variation for heritability. bioRxiv 2014:006387.

24. Beerenwinkel N, Günthard HF, Roth V, Metzner KJ: Challenges and opportunities in estimating viral genetic diversity from next-generation sequencing data. Front Microbiol 2012, 3:329.

25. Li H: Aligning sequence reads, clone sequences and assembly contigs with BWA-MEM. arXiv:13033997 2013:3.

26. Zook JM, Chapman B, Wang J, Mittelman D, Hofmann O, Hide W, Salit M: Integrating human sequence data sets provides a resource of benchmark SNP and indel genotype calls. Nat Biotechnol 2014, 32:246–251.

27. Giallonardo F Di, Töpfer A, Rey M, Prabhakaran S, Duport Y, Leemann C, Schmutz S, Campbell NK, Joos B, Lecca MR, Patrignani A, Däumer M, Beisel C, Rusert P, Trkola A, Günthard HF, Roth V, Beerenwinkel N, Metzner KJ: Full-length haplotype reconstruction to infer the structure of heterogeneous virus populations. Nucleic Acids Res 2014:gku537–.

28. Prabhakaran S, Rey M, Zagordi O, Beerenwinkel N, Roth V: HIV-Haplotype Inference using a Constraint-based Dirichlet Process Mixture Model. In NIPS Work Mach Learn Comput Biol; 2010:1–4.

29. Do CB, Mahabhashyam MSP, Brudno M, Batzoglou S: ProbCons: Probabilistic consistency. based multiple sequence alignment. Genome Res 2005, 15:330–40.

30. Sarawagi S, Cohen WW: Semi-Markov conditional random fields for information extraction. In Adv Neural Inf Process Syst; 2004.

31. Chin CHS, Alexander DH, Marks P, Klammer AA, Drake J, Heiner C, Clum A, Copeland A, Huddleston J, Eichler EE, Turner SW, Korlach J: Nonhybrid, finished microbial genome assemblies from long-read SMRT sequencing data. Nat Methods 2013, 10:563–9.

